## supplementary information for "Metabolic Derangement in Polycystic Kidney Disease Mouse Models Is Ameliorated by Mitochondrial-Targeted Antioxidants"

**Supplementary Figures:**
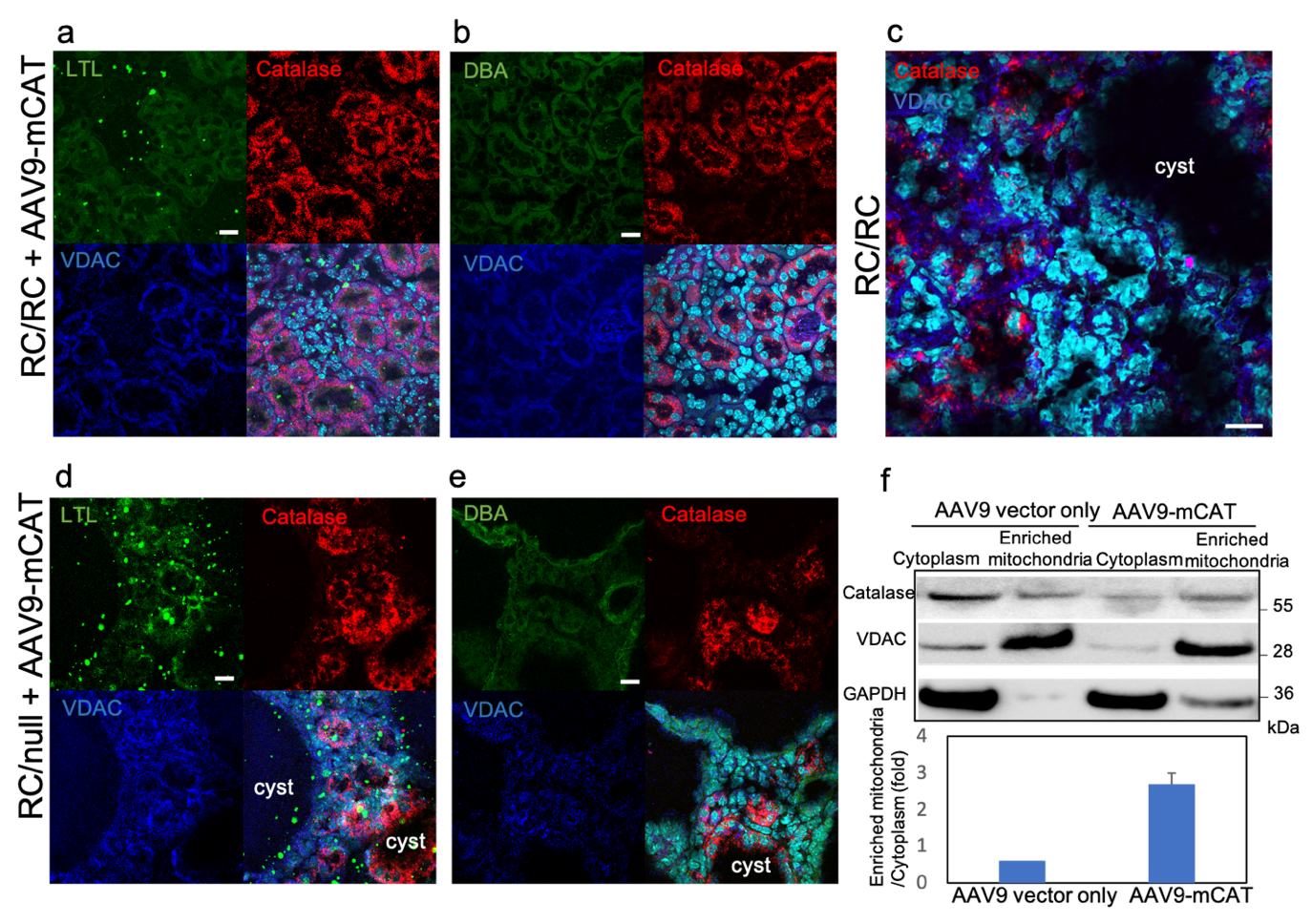

**Supplementary Figure 1.** co-immunostaining of frozen kidney sections of (a-c) RC/RC and RC/RC treated with AAV9-mCAT and (d-e) RC/null treated with AAV9-mCAT with specific proximal or distal tubular markers (LTL and DBA, respectively), VDAC (mitochondrial marker) and catalase (Scale bars:10um). (f) Western-blot of catalase in cytoplasm and enriched mitochondrial of RC/RC and RC/RC+AAV9-mCAT mice; n=3 per group

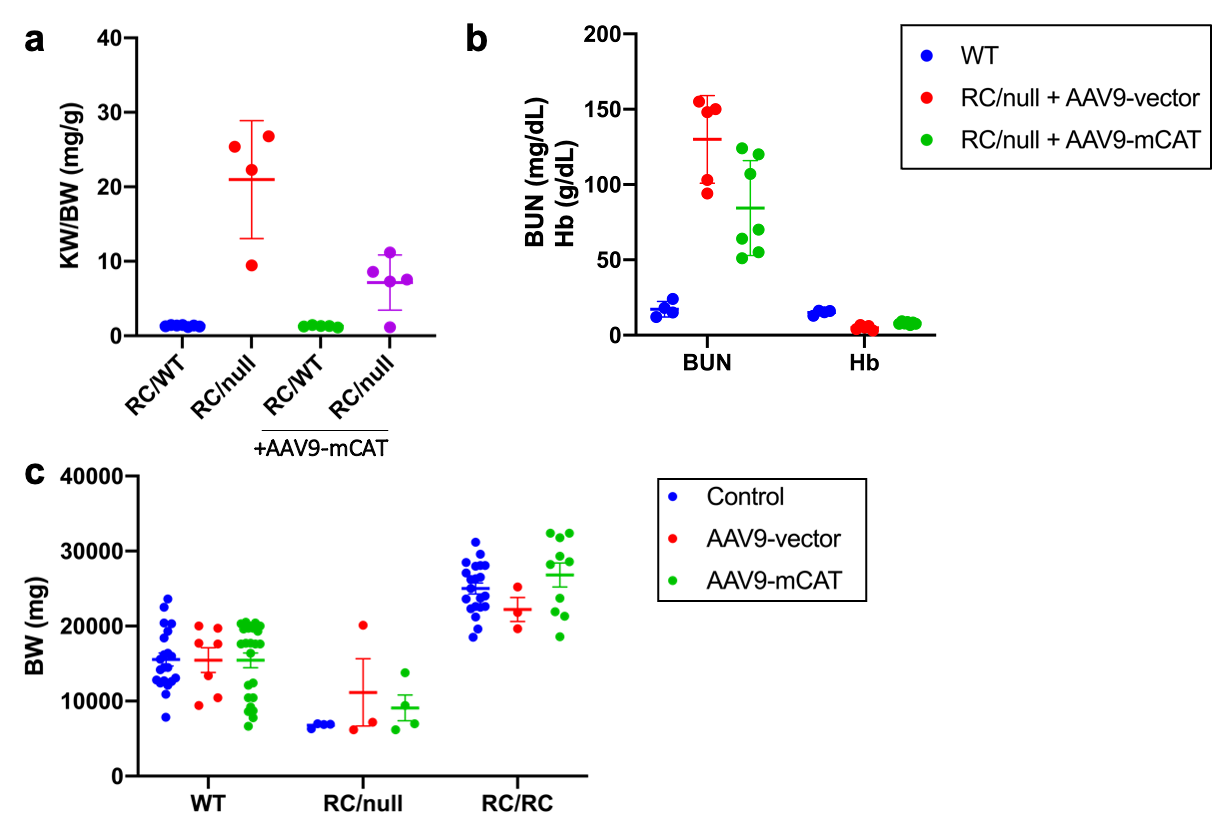

**Supplementary Figure 2.** (a) Individual data points of KW/BW in RC/WT and RC/null mice with or without treatment with AAV9-mCAT. (b) BUN (mg/dL) and Hb (g/dL) individual data points of the indicated genotypes and treatment groups (n=4-8). (c) Effect of treatment with AAV9-vector or AAV9-mCAT on body weight of RC/RC, RC/null and RC/WT mice (n=4-30).

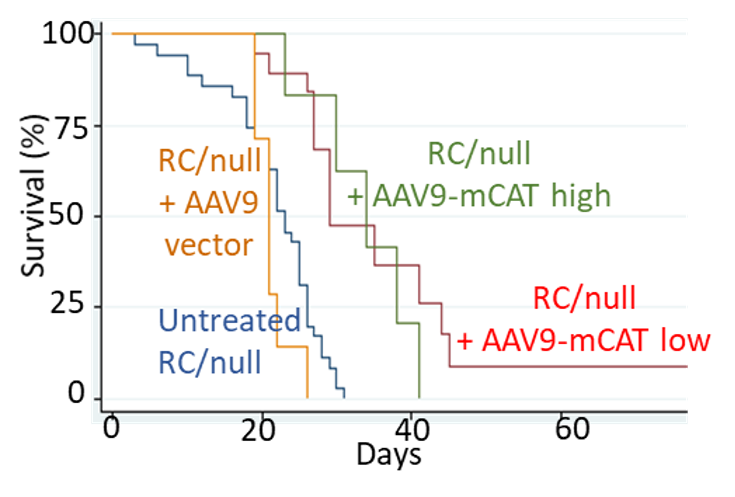

**Supplementary Figure 3.** Survival curves of RC/null mice or RC/null mice treated with AAV-9 mCAT, including both low (2*10^9^/g) and high (10^10^/g) doses; n=31 for RC/null, n=19 for low dose and 9 for high dose mCAT. p values determined by log-rank test; p<0.001.

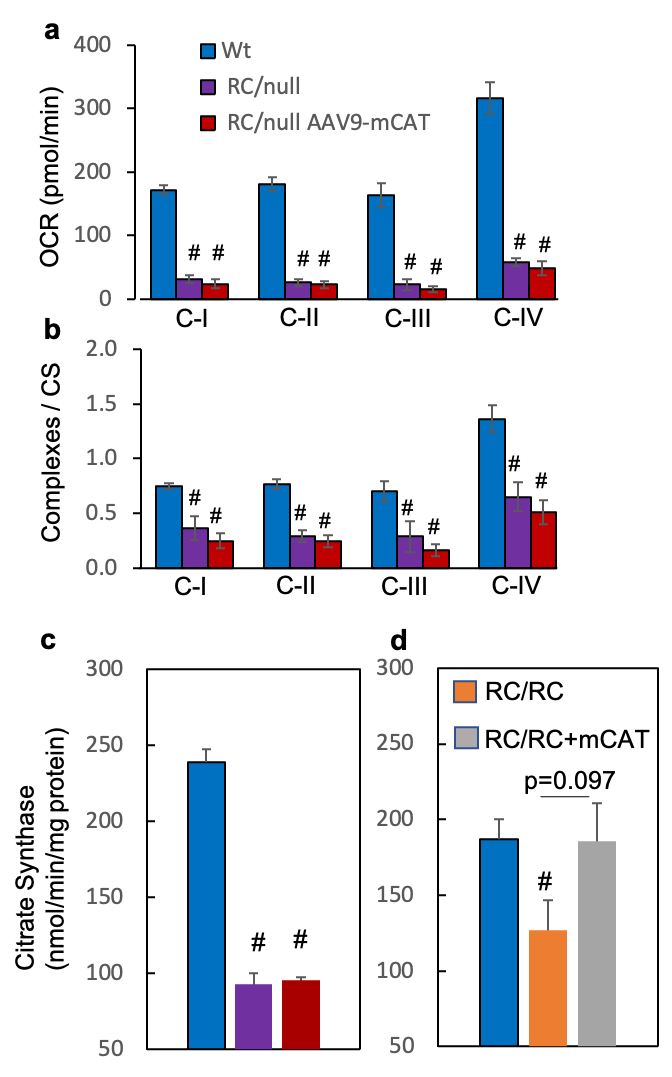

**Supplementary Figure 4.** Mitochondrial complex activity in kidney lysates from 5 WT, 4 RC/null mice and 4 RC/null mice treated with mCAT (RC/null AAV9-mCAT), as measured by Seahorse analyzer, (a) before and (b) after values were normalized to citrate synthase (CS) activity; (c) CS activity. n=4-5 each group; (d) CS activity in a separate experiment in RC/RC kidney lysates, N=6 each group; #p<0.05 compared with WT (blue bar).

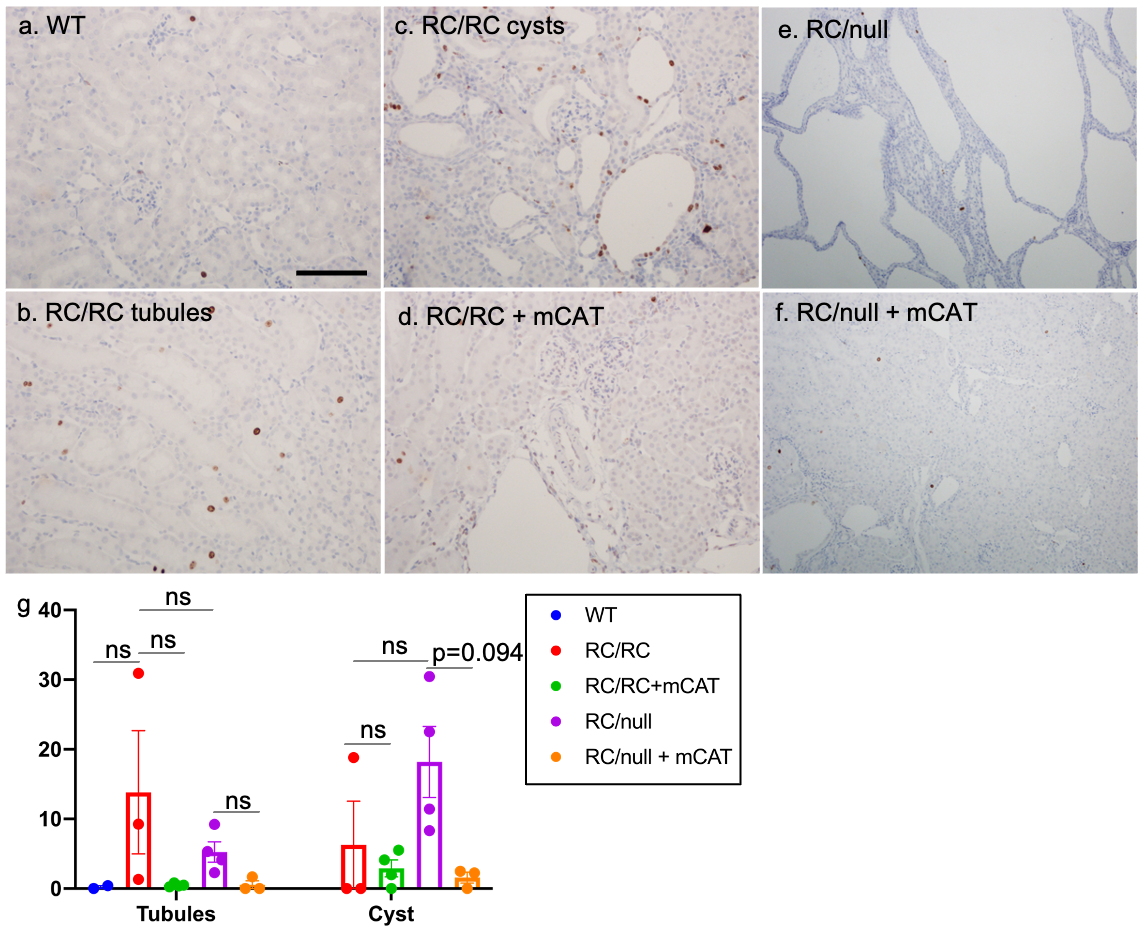

**Supplementary Figure 5.** (a-f) Representative images of Ki-67 staining of tubules and cyst-lining epithelium in kidneys of WT, RC/RC and RC/null mice untreated or treated with AAV9-mCAT. (Scale bar:100um) (g) Individual data points of indicated genotypes and groups presented in g-h; multiple images were taken from kidney tissues of 3-4 mice per group.
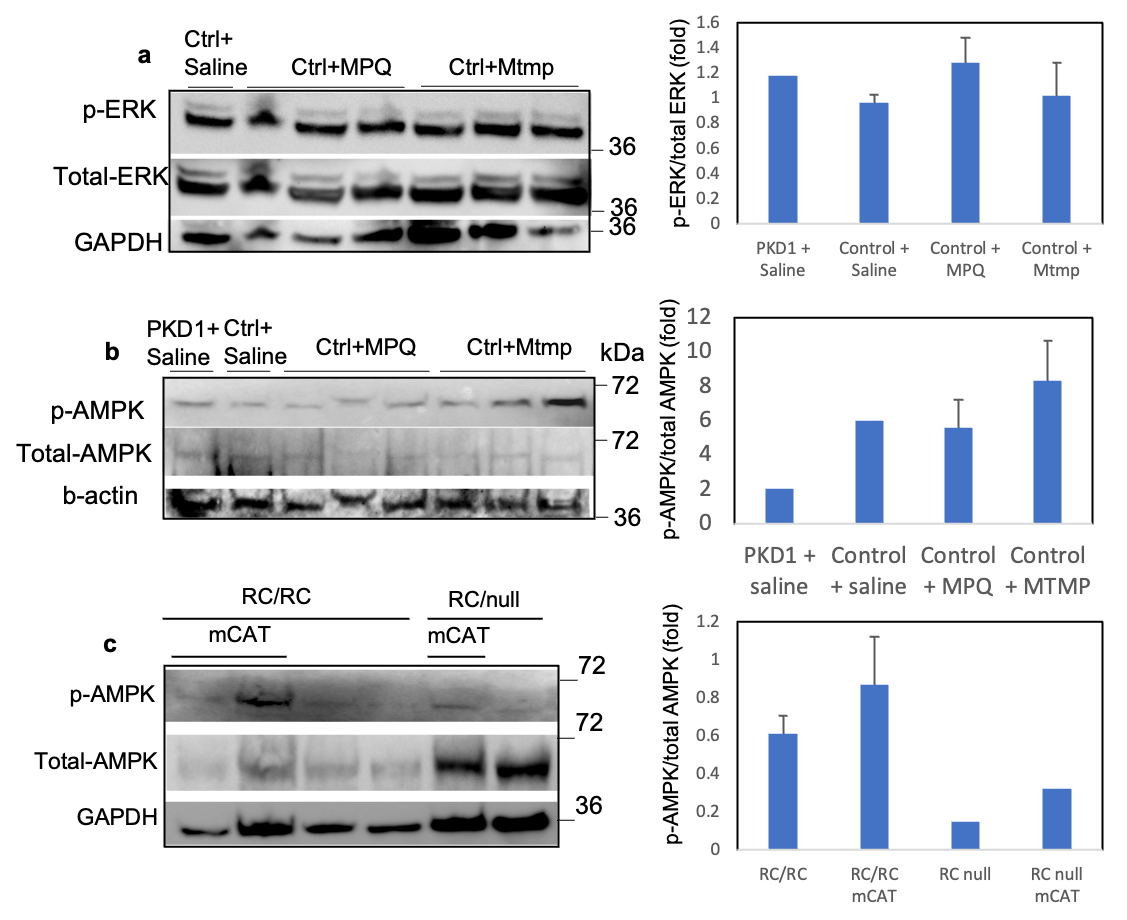

**Supplementary Figure 6.** (a-b) The effect of MPQ and Mtmp treatment on the phosphorylation of AMPK and ERK shown by immunoblotting of (a) p-ERK and (b) p-AMPK in HK2 control cells compared with PKD1 mutant cells and quantification of indicated groups. (c) AMPK phosphorylation in mouse kidneys of RC/RC, RC/null and RC/RC, RC/null treated with AAV9-mCAT and quantification. GAPDH and b-actin were used as loading control; n=3

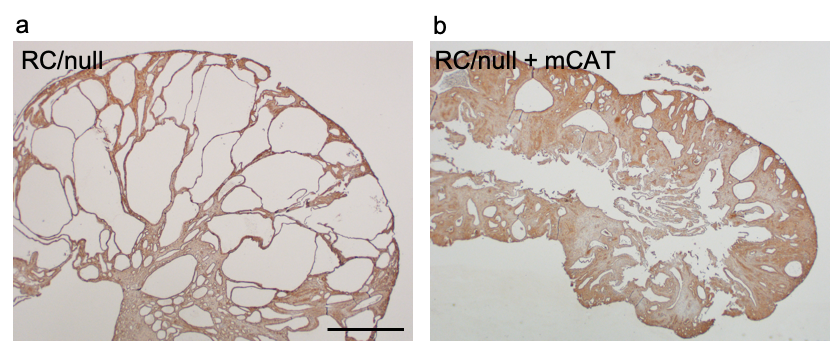

**Supplementary Figure 7.** (a) IHC staining for nitrotyrosine in RC/null and (b) RC/null mice treated with AAV9-mCAT; scale bar: 1mm; n=3 per group

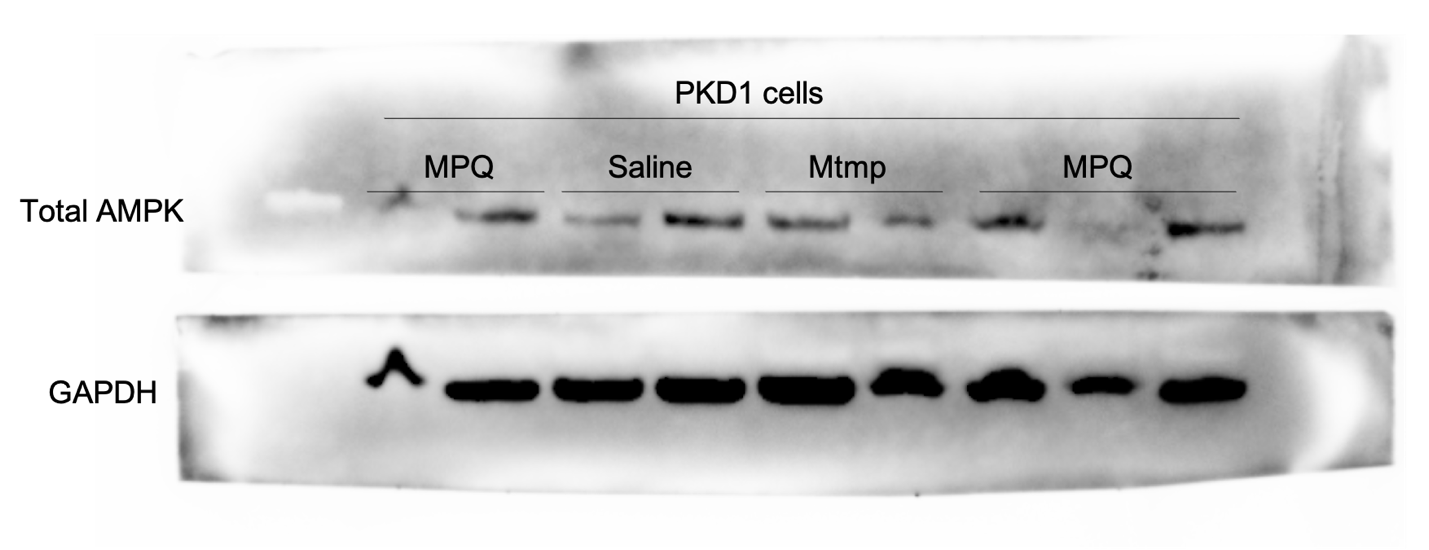

**Supplementary Figure 8.** Uncropped blot of Figure 6c. Immunoblotting of AMPK in PKD1 mutant cells (WT9-7). Mtmp: Mito-Tempo, MPQ: mito-paraquat.

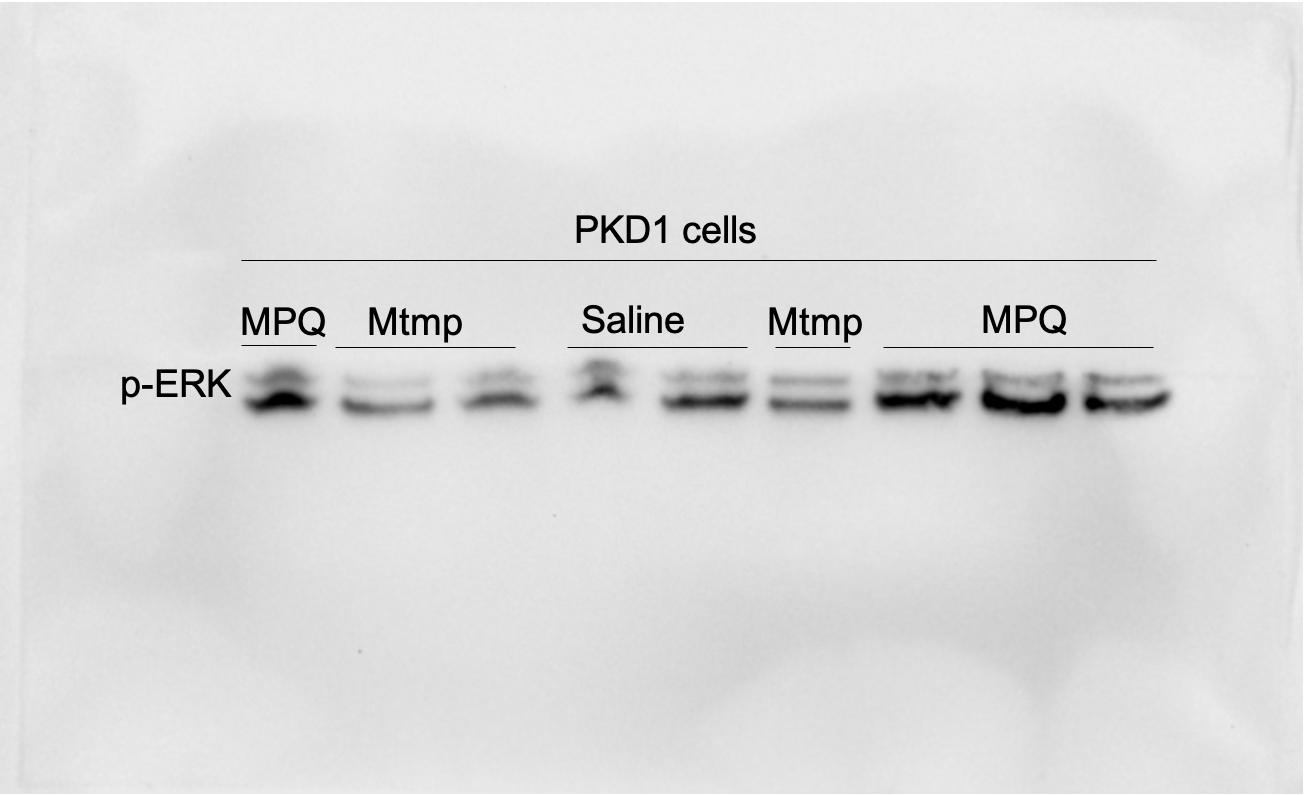

**Supplementary Figure 9.** Uncropped blot of Figure 6d. Immunoblotting of p-ERK in PKD1 mutant cells (WT9-7). Mtmp: Mito-Tempo, MPQ: mito-paraquat.

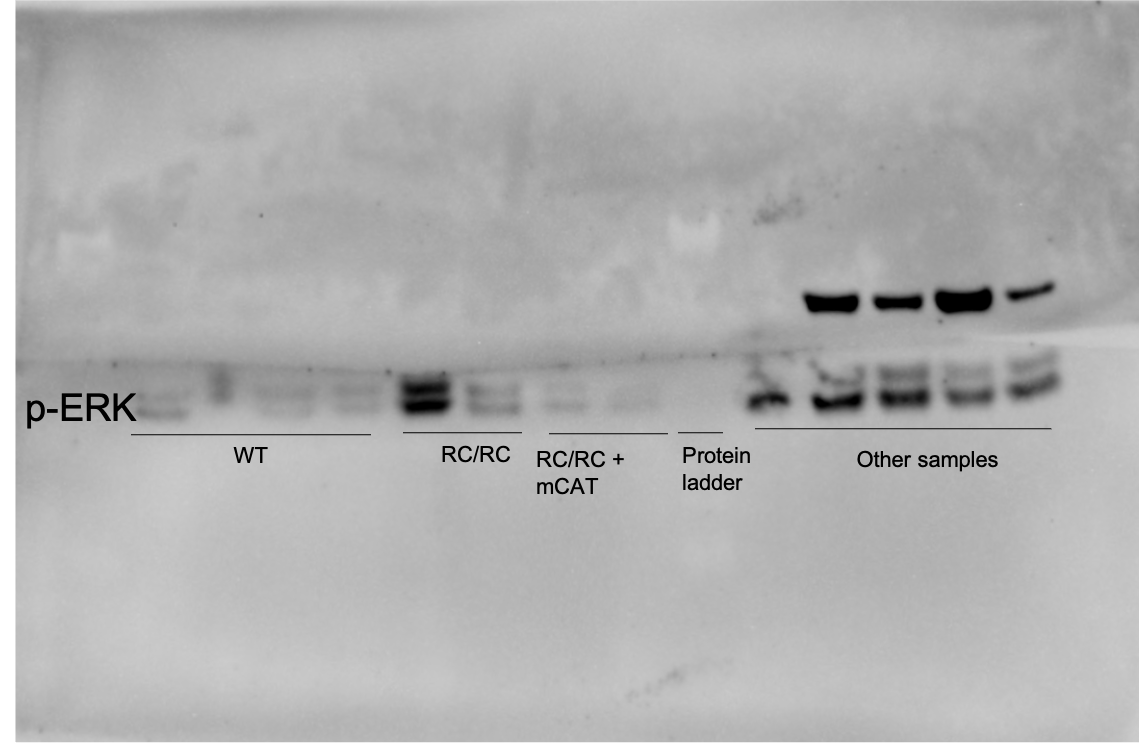

**Supplementary Figure 10.** Uncropped blot of Figure 6g. Immunoblotting of p-ERK in WT, RC/RC and RC/RC treated with AAV9-mCAT

**Supplementary Methods:**

**Immunofluorescence**

Frozen sections (5 μm) were fixed in 4% paraformaldehyde for 10 minutes, washed and permeabilized with 0.05% Triton-x, washed then blocked with normal donkey or goat serum for 1 hour, followed by staining with primary antibodies: goat anti-Catalase (R&D AF-3398-SP, 1:10) and rabbit anti VDAC-1 (Abcam 15895, 1:100) at 4ºC overnight. After multiple washings, sections were incubated with secondary antibody AlexaFluor 555 donkey anti goat (1:200, BD Biosciences) and AlexaFluo 488 goat anti rabbit (1:200, BD Biosciences) or donkey-anti-rabbit IgG Alexa Fluor 647 (1:200, Invitrogen A-31573) and Dolichos Biflorus Agglutinin (DBA), Fluorescein (Vector lab, FL-1031, 1:50) or Lotus Tetragonolobus Lectin (LTL), Fluorescein (Vector labs, FL-1321-2, 1:50) for 90 minutes then washed and counterstained with Hoechst 33342 for 5 minutes. The images were taken using a Leica confocal SP8.

**Western Blot**

Kidney tissues were homogenized in a glass homogenizer in 0.3 ml RIPA buffer with protease and phosphatase inhibitors (Roche). Homogenates were sonicated, and centrifuged at 10,000xg for 20 minutes and supernatants were collected. For cell cultures, cells were washed with ice-cold PBS after discarding the medium. Then, PBS was discarded and 80-100ul of ice-cold RIPA lysis buffer was added and cells were scraped using a plastic cell-scraper and collected in a microcentrifuge tubes. Protein concentrations were determined using the BCA method (Pierce). Samples were boiled for 5 minutes prior to loading. Equal amounts of protein were loaded onto a NuPAGE 4-12% Bis-Tris gel (Novex). Electrophoresis was performed at 4^0^C in a MiniGel tank (Invitrogen) at 120V for 2 hours. Proteins were blotted at 4^0^C onto a PVDF membrane (Bio-Rad) using the same tank and a blot module (Invitrogen), at 20V for 5 hours. Blots were stained with Ponceau S to confirm equal loading and were blocked in 5% non-fat milk in PBS-Tween 20 (0.05%) (PBS-T). Blots were then incubated with primary antibodies (1:500 in 10% BSA) at 4^0^C for overnight (~12 hours), washed in PBS-T, and incubated with HRP-conjugated secondary antibody (1:10,000 in 5% milk-PBS-T) at room temperature for 1 hour. Blots were washed with PBS-T, exposed to West Femto chemiluminescent reagent, and imaged in a Bio-Rad Imager. Bands were quantified using Image J (NIH).

**Measurement of oxidative damage and redox status**

The levels of F2-isoprostanes in kidneys were determined^1^ with minor modifications. Briefly, 100 mg of tissue was homogenized in 10 ml of ice-cold Folch solution (CHCl3: MeOH, 2:1) containing butylated hydroxytoluene (BHT). The mixture was incubated at room temperature for 30 minutes. Two ml of 0.9% NaCl was added and mixed well. The homogenate was centrifuged at 3,000g for 5 minutes at 4 °C. The aqueous layer was discarded while the organic layer was secured and evaporated to dryness under N2 at 37 °C. The level of F2-isoprostanes in the tissues was expressed as nanograms of 8-Iso-PGF2α, per gram of tissue.

**Targeted Proteomics Analysis of Frozen Kidneys**

Twenty-seven kidneys (including WT, RC/RC, RC/RC + mCAT, RC/null and RC/null + mCAT) were frozen in liquid nitrogen, then homogenized in RIPA buffer containing protease inhibitor cocktail. One hundred µg of total protein from each sample was used for targeted proteomics analysis. Total proteins were mixed with 200 µL 1% SDS, 20 µL of BSA internal standard, heated for 15 minutes, and then precipitated with 1 mL acetone. The dried protein pellet was reconstituted in 100 µL Laemmli sample buffer and 20 µL (20 µg) was used to run a short (1.5 cm) SDS-PAGE gel. The gels were fixed and stained. Each sample was cut from the gel as the entire lane and divided into smaller pieces. The gel pieces were washed to remove Coomassie blue, reduced, alkylated, and digested overnight with trypsin. The mixture of peptides was extracted from the gel, evaporated to dryness in a SpeedVac and reconstituted in 150 µL of 1% acetic acid for analysis.

The analyses were carried out on a TSQ Quantiva triple quadrupole mass spectrometry system. The HPLC was an Ultimate 3000 nanoflow system with a 10 cm x 75 µm i.d. C18 reversed phase capillary column. 5 µL aliquots were injected and peptides were eluted with a 60-minute gradient of acetonitrile in 0.1% formic acid.

The mass spectrometer was operated in the selected reaction monitoring mode. For each protein, the method was developed to measure 2 ideal peptides. Assays for multiple proteins were bundled together in larger panels. Data were analyzed using the program Skyline to determine the integrated peak area of the appropriate chromatographic peaks. The response for each protein was calculated as the geometric mean of the peptide areas. These values were normalized to the response for the BSA standard. The samples were also analyzed on a Thermo QEx system in the LC-full scan MS mode. The total ion current in those analyses is an indication of the amount of material present in the sample for normalization.

Additional ‘universal detection’ runs, high resolution accurate mass (HRAM) were also performed using an orbitrap system (ThermoScientific QEx plus), as an additional type of data that could be re-interrogated when needed.

**Statistical Analysis of Targeted Proteomics Data**

The targeted proteomics data are analyzed pathway by pathway. Let $\mathcal{P}$ be the set of all proteins in a certain pathway. We consider the following model,

$$Y=\beta_{0}+\sum_{p\in\mathcal{P}} \left( \beta_{p}I_{p}+\tau_{p\_PKD1}(I_{p_{1}}+I_{p_{2}})+\tau_{p\_dose}I_{p_{2}}+\tau_{p\_mCAT}I_{p_{3}}\times(I_{p_{1}}+I_{p_{2}})+\tau_{p\_mCAT\_dose}I_{p_{3}}\times I_{p_{2}} \right)+\epsilon,$$

where *Y* is the log-transformed data, and $\beta_{0}$*,* $\beta_{p}$*,* $\tau_{p\_PKD1}$*,* $\tau_{p\_dose}$*,* $\tau_{p\_mCAT}$*,* $\tau_{p\_mCAT\_dose}$ are the overall mean, protein-specific effect relative to the overall mean, protein-specific effect of PKD1, protein-specific dose effect, protein-specific mCAT treatment effect, and protein-specific interaction effects of mCAT and dose, respectively. For a certain protein $p\in\mathcal{P}$, the indicator variables are defined as,

$$I_{p}=\left\{ \begin{aligned} 1, &observation from protein p, \\ 0, &otherwise \end{aligned} \right.$$

$$I_{p_{1}}=\left\{ \begin{aligned} 1, &observation from&&& protein p \mathrm{and} &&from RC/RC, \\ 0, &otherwise \end{aligned} \right.$$

$$I_{p_{2}}=\left\{ \begin{aligned} 1, &observation from&&& protein p \mathrm{and} &&from Rcnull, \\ 0, &otherwise \end{aligned} \right.$$

$$I_{p_{3}}=\left\{ \begin{aligned} 1, &observation from&&& protein p \mathrm{and} &&treated with mCAT, \\ 0, &otherwise \end{aligned} \right.$$

The parameters are grouped by proteins. For a given protein$p$, its group parameter vector is defined as$\boldsymbol{\beta}_{p}=(\beta_{p}, \tau_{p\_PKD1}, \tau_{p\_dose}, \tau_{p\_mCAT}, \tau_{p\_mCAT\_dose})^{T}\triangleq(\beta_{p1},\beta_{p2},\beta_{p3},\beta_{p4},\beta_{p5})$. The overall parameter vector for the model is$\boldsymbol{\beta}=\left( {\beta_{0}\boldsymbol{,\beta}}_{1}, \ldots, \boldsymbol{\beta}_{G} \right)^{T}$, where *G* is the number of groups in the pathway. We selected important groups and important members within groups simultaneously. This is referred to as bi-level selection by Breheny & Huang^2^. They also proposed the composite Minimax Concave Penalty (cMCP) to implement bi-level selection. Let $f_{\lambda,a}$be the minimax concave penalty function defined by,

$f_{\lambda,a}\left( \theta\right)=\left\{ \begin{aligned} \lambda\theta-\frac{\theta^{2}}{2a}, if \theta\leq a\lambda\\ \frac{1}{2}a\lambda^{2}, if \theta>a\lambda\end{aligned} \right.$ ,

where $\lambda$ is the regularization parameter and *a* is the tuning parameter. In our model, the cMCP minimizes,

$$Q\left( \boldsymbol{\beta} \right)=\frac{1}{2n}\left\| Y-X\boldsymbol{\beta} \right\|^{2}+\sum_{g=1}^{G} f_{O}\left( \sum_{k=1}^{K_{g}} f_{I}\left( |\beta_{gk}| \right) \right) ,$$

where *X* is the design matrix and $K_{g}=5$ is the number of parameters in each group. Here $f_{O}=f_{\lambda,b}$ and $f_{I}=f_{\lambda,a}$ are called outer and inner penalty functions, respectively. We follow Breheny & Huang (2009) in setting $a=3$ and $b=K_{g}a\lambda/2$. The regularization parameter $\lambda$ is determined by 10-fold cross validation and then the best subset of parameters is selected, using the R package grpreg. The estimates are derived by fitting unpenalized linear regression models with the selected variables, i.e., whose coefficients are deemed non-zero if the penalized estimates are non-zero.
