## supplementary data 1 for "Metabolic Derangement in Polycystic Kidney Disease Mouse Models Is Ameliorated by Mitochondrial-Targeted Antioxidants"

| Protein ID | Protein name | Targeted Proteome Measurement of Abundance (pmol/100µg total protein) | | | | | | | | | | Magnitude of Effect (%)* | | | |
|---|---|---|---|---|---|---|---|---|---|---|---|---|---|---|---|
| | | WT | | RC/RC | | RC/RC+mCAT | | RC/null | | RC/null+mCAT | | PKD1 mutation effect | mCAT effect | PKD1 dose effect | PKD1 dose & mCAT interaction |
| | | Mean | SE | Mean | SE | Mean | SE | Mean | SE | Mean | SE | | | | |
| **Fatty Acid Beta Oxidation (FAO)** | | | | | | | | | | | | | | | |
| **Fabp3** | fatty acid binding protein 3 | 0.384 | 0.028 | 0.234 | 0.016 | 0.441 | 0.124 | 0.160 | 0.041 | 0.157 | 0.007 | -0.398 | 0.630 | -0.416 | -0.281 |
| **Fabp4** | fatty acid binding protein 4 | 4.402 | 1.165 | 2.848 | 0.723 | 2.431 | 0.529 | 1.793 | 0.820 | 1.473 | 0.527 | -0.448 | -0.027 | -0.381 | -0.092 |
| **Cpt1a** | carnitine palmitoyltransferase 1a | 1.043 | 0.091 | 1.115 | 0.124 | 1.322 | 0.125 | 0.407 | 0.026 | 0.578 | 0.050 | N.S | 0.227 | -0.613 | 0.154 |
| **Cpt1b** | carnitine palmitoyltransferase 1b | 0.170 | 0.019 | 0.092 | 0.010 | 0.140 | 0.019 | 0.028 | 0.008 | 0.058 | 0.020 | -0.470 | 0.520 | -0.727 | 0.386 |
| **Cpt2** | carnitine palmitoyltransferase 2 | 0.268 | 0.029 | 0.200 | 0.025 | 0.208 | 0.022 | 0.072 | 0.016 | 0.097 | 0.006 | -0.299 | 0.091 | -0.637 | 0.322 |
| **Hadh** | hydroxyacyl-Coenzyme A dehydrogenase(trifunctional protein) | 2.987 | 0.445 | 2.758 | 0.460 | 2.990 | 0.366 | 0.681 | 0.086 | 0.776 | 0.003 | -0.180 | 0.189 | -0.723 | N.S |
| **Hadha** | hydroxyacyl-CoA dehydrogenase/3-ketoacyl-CoA thiolase/enoyl-CoA hydratase | 1.747 | 0.594 | 0.632 | 0.042 | 0.802 | 0.048 | 0.349 | 0.034 | 0.485 | 0.038 | -0.595 | 0.316 | -0.423 | N.S |
| **Hadhb** | (trifunctional protein), alpha and beta subunit | 1.262 | 0.319 | 0.535 | 0.032 | 0.604 | 0.051 | 0.299 | 0.049 | 0.363 | 0.030 | -0.544 | 0.125 | -0.452 | 0.110 |
| **acsl1** | acyl-CoA synthetase long-chain family member 1 | 2.073 | 0.302 | 0.857 | 0.064 | 1.146 | 0.126 | 0.435 | 0.043 | 0.595 | 0.093 | -0.583 | 0.333 | -0.484 | N.S |
| **Acad11** | acyl-Coenzyme A dehydrogenase family, member 11 | 0.413 | 0.088 | 0.391 | 0.051 | 0.475 | 0.049 | 0.092 | 0.023 | 0.129 | 0.041 | N.S | 0.262 | -0.770 | 0.104 |
| **Acadvl** | acyl-Coenzyme A dehydrogenase, very long chain | 0.973 | 0.263 | 0.505 | 0.050 | 0.645 | 0.029 | 0.220 | 0.016 | 0.326 | 0.081 | -0.454 | 0.354 | -0.534 | N.S |
| **Acadl** | acyl-Coenzyme A dehydrogenase, long-chain | 1.533 | 0.394 | 1.091 | 0.096 | 1.408 | 0.059 | 0.515 | 0.034 | 0.712 | 0.172 | -0.245 | 0.327 | -0.516 | N.S |
| **Acadm** | acyl-Coenzyme A dehydrogenase, medium chain | 3.655 | 0.731 | 2.641 | 0.348 | 3.010 | 0.155 | 0.533 | 0.062 | 0.617 | 0.073 | -0.298 | 0.216 | -0.789 | N.S |
| **Acads** | acyl-Coenzyme A dehydrogenase, short chain | 0.469 | 0.068 | 0.633 | 0.089 | 0.662 | 0.049 | 0.172 | 0.011 | 0.173 | 0.011 | 0.194 | 0.098 | -0.716 | N.S |
| **Ehhadh** | enoyl-Coenzyme A, hydratase/3-hydroxyacyl Coenzyme A dehydrogenase | 2.089 | 0.640 | 1.563 | 0.203 | 2.073 | 0.271 | 0.197 | 0.025 | 0.349 | 0.115 | -0.168 | 0.388 | -0.866 | 0.191 |
| **echs1** | enoyl Coenzyme A hydratase, short chain, 1, mitochondrial | 0.580 | 0.125 | 0.490 | 0.057 | 0.562 | 0.060 | 0.173 | 0.033 | 0.194 | 0.019 | N.S | 0.129 | -0.656 | N.S |
| **Eci1** | enoyl-Coenzyme A delta isomerase 1 | 0.561 | 0.094 | 0.469 | 0.057 | 0.532 | 0.049 | 0.205 | 0.028 | 0.218 | 0.016 | -0.090 | 0.159 | -0.562 | N.S |
| **Eci2** | enoyl-Coenzyme A delta isomerase 2 | 0.075 | 0.015 | 0.059 | 0.006 | 0.064 | 0.003 | 0.036 | 0.004 | 0.044 | 0.008 | -0.199 | 0.138 | -0.358 | N.S |
| **Acaa1a/b** | acetyl-Coenzyme A acyltransferase 1A/1B | 0.140 | 0.024 | 0.101 | 0.012 | 0.130 | 0.011 | 0.062 | 0.011 | 0.075 | 0.008 | -0.282 | 0.313 | -0.400 | N.S |
| **Acaa2** | acetyl-Coenzyme A acyltransferase 2 (mitochondrial 3-oxoacyl-CoA thiolase) | 2.488 | 0.426 | 0.983 | 0.075 | 1.284 | 0.091 | 0.537 | 0.029 | 0.735 | 0.080 | -0.601 | 0.335 | -0.434 | N.S |
| **crat** | carnitine acetyltransferase | 0.198 | 0.058 | 0.109 | 0.011 | 0.126 | 0.008 | 0.044 | 0.005 | 0.056 | 0.005 | -0.408 | 0.189 | -0.587 | 0.085 |
| **gpi** | glucose phosphate isomerase 1 | 0.243 | 0.020 | 0.166 | 0.024 | 0.143 | 0.022 | 0.170 | 0.030 | 0.240 | 0.003 | -0.364 | -0.120 | 0.064 | 0.678 |
| **FA biosynthesis and ketogenesis** | | | | | | | | | | | | | | | |
| **decr1** | 2,4-dienoyl CoA reductase 1, mitochondrial | 1.313 | 0.177 | 1.160 | 0.136 | 1.247 | 0.133 | 0.425 | 0.057 | 0.513 | 0.037 | N.S | 0.052 | -0.639 | 0.175 |
| **Acot13** | acyl-CoA thioesterase 13 | 1.595 | 0.133 | 1.111 | 0.117 | 1.735 | 0.253 | 0.471 | 0.009 | 0.565 | 0.057 | -0.326 | 0.535 | -0.558 | -0.226 |
| **Hsd17b4** | hydroxysteroid (17-beta) dehydrogenase 4 | 1.001 | 0.195 | 1.006 | 0.131 | 1.047 | 0.118 | 0.440 | 0.055 | 0.468 | 0.040 | N.S | 0.079 | -0.540 | N.S |
| **Hmgcs1** | 3-hydroxy-3-methylglutaryl-Coenzyme A synthase 1 | 0.071 | 0.007 | 0.105 | 0.013 | 0.119 | 0.015 | 0.072 | 0.008 | 0.070 | 0.002 | 0.394 | 0.156 | -0.269 | -0.152 |
| **Hmgcs2** | 3-hydroxy-3-methylglutaryl-Coenzyme A synthase 2 | 0.370 | 0.117 | 0.149 | 0.017 | 0.235 | 0.032 | 0.240 | 0.032 | 0.224 | 0.033 | -0.553 | 0.562 | 0.646 | -0.399 |
| **HmgcL** | 3-hydroxy-3-methylglutaryl-Coenzyme A lyase | 0.879 | 0.147 | 0.866 | 0.119 | 1.019 | 0.123 | 0.306 | 0.026 | 0.302 | 0.028 | N.S | 0.209 | -0.624 | -0.183 |
| **Bdh1** | 3-hydroxybutyrate dehydrogenase, type 1 | 1.791 | 0.437 | 2.062 | 0.212 | 2.436 | 0.286 | 0.297 | 0.037 | 0.351 | 0.019 | 0.223 | 0.191 | -0.852 | N.S |
| **Glycolysis and Gluconeogenesis** | | | | | | | | | | | | | | | |
| **pygm** | muscle glycogen phosphorylase | 0.025 | 0.003 | 0.014 | 0.002 | 0.031 | 0.011 | 0.010 | 0.004 | 0.033 | 0.010 | -0.498 | 1.039 | -0.291 | 0.670 |
| **pygb** | brain glycogen phosphorylase | 0.072 | 0.010 | 0.053 | 0.004 | 0.052 | 0.003 | 0.063 | 0.004 | 0.064 | 0.009 | -0.259 | N.S | 0.214 | N.S |
| **slc2a4** | solute carrier family 2 (facilitated glucose transporter), member 4 | 0.136 | 0.024 | 0.048 | 0.004 | 0.078 | 0.006 | 0.036 | 0.001 | 0.044 | 0.003 | -0.639 | 0.619 | -0.233 | -0.254 |
| **hk1** | hexokinase 1 | 0.531 | 0.097 | 0.477 | 0.077 | 0.405 | 0.021 | 0.508 | 0.089 | 0.406 | 0.054 | -0.109 | -0.113 | N.S | N.S |
| **tkt** | transketolase | 1.681 | 0.246 | 1.188 | 0.055 | 1.090 | 0.060 | 1.496 | 0.146 | 1.998 | 0.118 | -0.273 | -0.083 | 0.252 | 0.473 |
| **Taldo1** | transaldolase 1 | 0.025 | 0.004 | 0.021 | 0.002 | 0.018 | 0.001 | 0.022 | 0.005 | 0.019 | 0.001 | -0.163 | -0.103 | N.S | N.S |
| **gpi** | glucose phosphate isomerase 1 | 0.247 | 0.013 | 0.156 | 0.023 | 0.147 | 0.020 | 0.150 | 0.025 | 0.223 | 0.018 | -0.419 | N.S | N.S | 0.552 |
| **pfkl** | phosphofructokinase, liver, B-type | 0.191 | 0.028 | 0.163 | 0.014 | 0.179 | 0.018 | 0.219 | 0.022 | 0.162 | 0.011 | -0.106 | N.S | 0.310 | -0.252 |
| **pfkm** | phosphofructokinase, muscle | 0.085 | 0.003 | 0.065 | 0.005 | 0.093 | 0.008 | 0.050 | 0.007 | 0.053 | 0.006 | -0.251 | 0.429 | -0.236 | -0.255 |
| **aldoa** | aldolase A, fructose-bisphosphate | 1.439 | 0.328 | 0.995 | 0.051 | 1.004 | 0.054 | 1.198 | 0.162 | 1.076 | 0.054 | -0.265 | N.S | 0.137 | N.S |
| **aldob** | aldolase B, fructose-bisphosphate | 9.498 | 1.824 | 8.474 | 1.146 | 9.561 | 0.539 | 3.192 | 0.457 | 3.286 | 0.208 | -0.138 | 0.228 | -0.599 | -0.138 |
| **tpi** | triosephosphate isomerase 1 | 0.744 | 0.091 | 0.781 | 0.095 | 0.768 | 0.081 | 0.572 | 0.056 | 0.546 | 0.033 | N.S | N.S | -0.248 | N.S |
| **gapdh** | glyceraldehyde-3-phosphate dehydrogenase | 6.865 | 0.575 | 4.023 | 0.239 | 3.303 | 0.126 | 5.828 | 0.460 | 4.676 | 0.674 | -0.412 | -0.183 | 0.427 | N.S |
| **pgk1** | phosphoglycerate kinase 1 | 1.152 | 0.126 | 0.848 | 0.063 | 0.849 | 0.027 | 0.563 | 0.054 | 0.395 | 0.031 | -0.260 | N.S | -0.337 | -0.293 |
| **pgam2** | phosphoglycerate mutase 2 | 0.417 | 0.141 | 0.239 | 0.035 | 0.309 | 0.035 | 0.097 | 0.010 | 0.141 | 0.022 | -0.405 | 0.406 | -0.546 | N.S |
| **eno1** | enolase 1, alpha non-neuron | 3.997 | 0.489 | 2.583 | 0.202 | 2.241 | 0.128 | 2.050 | 0.208 | 2.096 | 0.214 | -0.368 | -0.079 | -0.145 | N.S |
| **pkm2** | pyruvate kinase, muscle | 2.313 | 0.247 | 1.311 | 0.100 | 1.353 | 0.044 | 1.939 | 0.216 | 1.798 | 0.145 | -0.424 | N.S | 0.417 | N.S |
| **ldha** | lactate dehydrogenase A | 5.600 | 0.505 | 6.022 | 0.618 | 6.073 | 0.835 | 4.897 | 0.403 | 4.050 | 0.108 | N.S | N.S | -0.149 | -0.165 |
| **ldhb** | lactate dehydrogenase B | 5.692 | 0.701 | 5.343 | 0.506 | 6.018 | 0.572 | 2.982 | 0.432 | 2.687 | 0.096 | N.S | 0.113 | -0.453 | -0.162 |
| **pc** | Pyruvate carboxylase | 3.225 | 0.479 | 2.817 | 0.428 | 3.265 | 0.409 | 0.695 | 0.153 | 1.163 | 0.245 | -0.213 | 0.248 | -0.738 | 0.384 |
| **mdh1** | malate dehydrogenase 1, NAD (soluble) | 8.092 | 0.699 | 6.472 | 0.872 | 7.572 | 0.750 | 2.793 | 0.145 | 3.608 | 0.635 | -0.253 | 0.233 | -0.531 | N.S |
| **hspd1** | heat shock protein 1 (chaperonin) | 1.486 | 0.190 | 1.236 | 0.110 | 1.041 | 0.096 | 0.685 | 0.098 | 0.863 | 0.048 | -0.178 | -0.148 | -0.443 | 0.523 |
| **Antioxidants** | | | | | | | | | | | | | | | |
| **akr1b1** | aldo-keto reductase family 1, member B3 (aldose reductase) | 1.357 | 0.222 | 0.410 | 0.100 | 0.552 | 0.097 | 0.290 | 0.013 | 0.273 | 0.007 | -0.728 | 0.423 | -0.174 | -0.339 |
| **aldh2** | aldehyde dehydrogenase 2, mitochondrial | 2.440 | 0.284 | 2.396 | 0.208 | 2.619 | 0.147 | 1.051 | 0.106 | 1.657 | 0.285 | N.S | 0.108 | -0.557 | 0.397 |
| **cat** | Catalase | 2.214 | 0.306 | 1.695 | 0.268 | 2.443 | 0.233 | 0.349 | 0.068 | 0.548 | 0.064 | -0.318 | 0.638 | -0.777 | N.S |
| **sod1** | superoxide dismutase 1, soluble | 1.193 | 0.212 | 0.942 | 0.126 | 1.275 | 0.151 | 0.419 | 0.040 | 0.505 | 0.121 | N.S | 0.266 | -0.583 | N.S |
| **sod2** | superoxide dismutase 2, mitochondrial | 0.382 | 0.060 | 0.309 | 0.046 | 0.398 | 0.029 | 0.097 | 0.015 | 0.136 | 0.036 | -0.231 | 0.378 | -0.674 | N.S |
| **gpi** | glucose phosphate isomerase 1 | 0.246 | 0.012 | 0.157 | 0.023 | 0.148 | 0.020 | 0.151 | 0.024 | 0.214 | 0.016 | -0.416 | N.S | N.S | 0.488 |
| **gpx1** | glutathione peroxidase 1 | 0.395 | 0.069 | 0.280 | 0.048 | 0.244 | 0.022 | 0.135 | 0.022 | 0.144 | 0.011 | -0.345 | N.S | -0.450 | N.S |
| **gpx4** | glutathione peroxidase 4 | 0.370 | 0.042 | 0.398 | 0.040 | 0.520 | 0.055 | 0.190 | 0.011 | 0.209 | 0.052 | N.S | 0.249 | -0.545 | N.S |
| **gsr** | glutathione reductase | 0.369 | 0.048 | 0.223 | 0.019 | 0.225 | 0.014 | 0.235 | 0.024 | 0.239 | 0.008 | -0.379 | N.S | N.S | N.S |
| **gsta3** | glutathione S-transferase, alpha 3 | 1.131 | 0.177 | 0.568 | 0.048 | 0.726 | 0.146 | 0.571 | 0.071 | 0.710 | 0.145 | -0.494 | 0.202 | N.S | N.S |
| **gstm1** | glutathione S-transferase, mu 1 | 4.889 | 0.921 | 3.992 | 0.426 | 4.238 | 0.456 | 3.261 | 0.458 | 3.065 | 0.564 | N.S | N.S | -0.219 | -0.063 |
| **gstp1** | glutathione S-transferase, pi 1 | 0.495 | 0.088 | 0.624 | 0.085 | 0.665 | 0.084 | 0.431 | 0.075 | 0.368 | 0.040 | N.S | N.S | -0.276 | -0.123 |
| **hspd1** | heat shock protein 1 (chaperonin) | 1.486 | 0.190 | 1.236 | 0.110 | 1.041 | 0.096 | 0.685 | 0.098 | 0.837 | 0.022 | N.S | N.S | -0.374 | N.S |
| **mdh1** | malate dehydrogenase 1, NAD (soluble) | 8.342 | 0.578 | 6.472 | 0.872 | 7.572 | 0.750 | 2.793 | 0.145 | 3.590 | 0.617 | -0.278 | 0.232 | -0.531 | N.S |
| **msra** | methionine sulfoxide reductase A | 1.005 | 0.151 | 0.864 | 0.111 | 0.994 | 0.123 | 0.297 | 0.062 | 0.265 | 0.007 | N.S | 0.076 | -0.695 | N.S |
| **phb** | prohibitin | 0.658 | 0.053 | 0.671 | 0.092 | 0.506 | 0.090 | 0.424 | 0.077 | 0.342 | 0.022 | N.S | -0.285 | -0.325 | N.S |
| **phb2** | prohibitin 2 | 0.571 | 0.100 | 0.470 | 0.083 | 0.272 | 0.038 | 0.274 | 0.042 | 0.218 | 0.015 | -0.253 | -0.334 | -0.290 | N.S |
| **prdx1** | peroxiredoxin 1 | 2.907 | 0.249 | 3.674 | 0.439 | 3.747 | 0.483 | 2.371 | 0.265 | 2.170 | 0.170 | N.S | N.S | -0.311 | -0.071 |
| **prdx2** | peroxiredoxin 2 | 0.659 | 0.050 | 0.669 | 0.043 | 0.722 | 0.074 | 0.594 | 0.027 | 0.558 | 0.032 | N.S | N.S | N.S | N.S |
| **prdx3** | peroxiredoxin 3 | 0.732 | 0.087 | 0.638 | 0.083 | 0.675 | 0.069 | 0.265 | 0.036 | 0.259 | 0.018 | N.S | N.S | -0.600 | N.S |
| **prdx5** | peroxiredoxin 5 | 4.485 | 1.005 | 3.569 | 0.684 | 4.191 | 0.610 | 1.219 | 0.049 | 0.891 | 0.052 | -0.267 | 0.289 | -0.606 | -0.433 |
| **prdx6** | peroxiredoxin 6 | 0.321 | 0.045 | 0.272 | 0.033 | 0.252 | 0.025 | 0.205 | 0.029 | 0.218 | 0.013 | -0.192 | N.S | -0.178 | N.S |
| **txn1** | thioredoxin 1 | 1.603 | 0.138 | 1.653 | 0.169 | 2.111 | 0.302 | 2.046 | 0.215 | 1.429 | 0.062 | N.S | 0.184 | N.S | -0.282 |
| **txnrd1** | thioredoxin reductase 1 | 0.256 | 0.021 | 0.226 | 0.014 | 0.255 | 0.017 | 0.216 | 0.011 | 0.195 | 0.021 | N.S | N.S | -0.097 | -0.103 |
| **TCA cycle and Related Proteins** | | | | | | | | | | | | | | | |
| **pdha1** | pyruvate dehydrogenase E1 alpha 1 | 1.630 | 0.345 | 1.099 | 0.112 | 1.171 | 0.070 | 0.423 | 0.049 | 0.457 | 0.005 | -0.314 | 0.105 | -0.604 | N.S |
| **pdhb** | pyruvate dehydrogenase (lipoamide) beta | 1.199 | 0.260 | 0.571 | 0.048 | 0.707 | 0.057 | 0.297 | 0.038 | 0.342 | 0.035 | -0.506 | 0.257 | -0.473 | -0.074 |
| **pdk1** | pyruvate dehydrogenase kinase, isoenzyme 1 | 0.019 | 0.004 | 0.012 | 0.001 | 0.011 | 0.001 | 0.009 | 0.001 | 0.011 | 0.001 | -0.349 | -0.066 | -0.291 | 0.364 |
| **pdk2** | pyruvate dehydrogenase kinase, isoenzyme2 | 0.073 | 0.011 | 0.048 | 0.004 | 0.044 | 0.004 | 0.022 | 0.003 | 0.023 | 0.003 | -0.348 | -0.048 | -0.530 | 0.112 |
| **dlat** | dihydrolipoamide S-acetyltransferase (E2 component of PDH) | 1.879 | 0.397 | 1.100 | 0.122 | 1.288 | 0.091 | 0.597 | 0.038 | 0.746 | 0.055 | -0.414 | 0.227 | -0.424 | N.S |
| **dld** | dihydrolipoamide dehydrogenase | 0.769 | 0.082 | 0.462 | 0.052 | 0.573 | 0.051 | 0.195 | 0.032 | 0.263 | 0.016 | -0.422 | 0.280 | -0.572 | 0.094 |
| **dlst** | dihydrolipoamide S-succinyltransferase (E2 comp of 2-oxo-glutarate complex) | 2.608 | 0.380 | 2.226 | 0.301 | 2.328 | 0.276 | 0.726 | 0.099 | 0.814 | 0.044 | -0.194 | 0.099 | -0.654 | 0.048 |
| **cs** | citrate synthase | 3.017 | 0.862 | 1.426 | 0.096 | 1.700 | 0.155 | 0.688 | 0.030 | 0.976 | 0.116 | -0.480 | 0.188 | -0.509 | 0.180 |
| **aco2** | aconitase 2, mitochondrial | 4.360 | 0.896 | 2.466 | 0.325 | 2.812 | 0.219 | 0.627 | 0.113 | 0.890 | 0.073 | -0.442 | 0.211 | -0.739 | 0.226 |
| **idh1** | isocitrate dehydrogenase 1 (NADP+), soluble | 3.783 | 0.316 | 2.429 | 0.187 | 2.847 | 0.143 | 1.626 | 0.093 | 1.379 | 0.080 | -0.369 | 0.195 | -0.316 | -0.289 |
| **idh2** | isocitrate dehydrogenase 2 (NADP+), mitochondrial | 5.181 | 1.048 | 3.146 | 0.264 | 3.822 | 0.258 | 1.201 | 0.138 | 1.497 | 0.188 | -0.377 | 0.242 | -0.611 | N.S |
| **idh3a** | isocitrate dehydrogenase 3 (NAD+) alpha | 0.560 | 0.085 | 0.393 | 0.035 | 0.395 | 0.020 | 0.178 | 0.006 | 0.228 | 0.011 | -0.299 | 0.036 | -0.531 | 0.240 |
| **idh3b** | isocitrate dehydrogenase 3 (NAD+) beta | 0.472 | 0.068 | 0.309 | 0.018 | 0.344 | 0.021 | 0.212 | 0.011 | 0.268 | 0.016 | -0.337 | 0.118 | -0.306 | 0.132 |
| **idh3g** | isocitrate dehydrogenase 3 (NAD+), gamma | 1.100 | 0.189 | 0.748 | 0.072 | 0.826 | 0.036 | 0.325 | 0.018 | 0.397 | 0.043 | -0.317 | 0.146 | -0.549 | 0.059 |
| **ogdh** | oxoglutarate (alpha-ketoglutarate) dehydrogenase (lipoamide) | 1.599 | 0.266 | 1.093 | 0.121 | 1.259 | 0.110 | 0.403 | 0.029 | 0.568 | 0.017 | -0.324 | 0.187 | -0.615 | 0.198 |
| **suclg1** | succinate-CoA ligase, GDP-forming, alpha subunit | 0.607 | 0.105 | 0.534 | 0.071 | 0.585 | 0.051 | 0.178 | 0.023 | 0.229 | 0.021 | -0.163 | 0.170 | -0.644 | 0.116 |
| **sucla2** | succinate-Coenzyme A ligase, ADP-forming, beta subunit | 1.067 | 0.144 | 0.717 | 0.060 | 0.780 | 0.037 | 0.321 | 0.023 | 0.365 | 0.042 | -0.332 | 0.121 | -0.539 | N.S |
| **sdha** | succinate dehydrogenase complex, subunit A, flavoprotein (Fp) | 2.519 | 0.254 | 2.030 | 0.248 | 2.567 | 0.129 | 0.635 | 0.034 | 0.768 | 0.065 | -0.235 | 0.343 | -0.667 | -0.102 |
| **sdhb** | succinate dehydrogenase complex, subunit B, iron sulfur (Ip) | 0.568 | 0.063 | 0.453 | 0.057 | 0.461 | 0.045 | 0.157 | 0.020 | 0.234 | 0.010 | -0.242 | 0.061 | -0.638 | 0.436 |
| **sdhc** | succinate dehydrogenase complex, subunit C, integral membrane protein | 0.220 | 0.019 | 0.201 | 0.026 | 0.265 | 0.033 | 0.100 | 0.002 | 0.099 | 0.006 | -0.138 | 0.350 | -0.466 | -0.274 |
| **fh1** | fumarate hydratase 1 | 0.741 | 0.121 | 0.585 | 0.060 | 0.629 | 0.028 | 0.225 | 0.027 | 0.249 | 0.022 | -0.217 | 0.120 | -0.605 | N.S |
| **mdh1** | malate dehydrogenase 1, NAD (soluble) | 7.830 | 0.500 | 6.555 | 0.800 | 7.529 | 0.744 | 2.882 | 0.152 | 3.514 | 0.643 | -0.212 | 0.190 | -0.533 | N.S |

| | | | | | | | | | | | | | | | |
|---|---|---|---|---|---|---|---|---|---|---|---|---|---|---|---|
| **mdh2** | malate dehydrogenase 2, NAD (mitochondrial) | 2.671 | 0.513 | 1.775 | 0.265 | 1.716 | 0.173 | 0.670 | 0.074 | 0.833 | 0.026 | -0.363 | 0.030 | -0.590 | 0.228 |
| **Glud1** | glutamate dehydrogenase 1 | 2.362 | 0.897 | 1.534 | 0.152 | 1.850 | 0.115 | 0.917 | 0.186 | 0.950 | 0.019 | N.S | 0.214 | -0.426 | -0.095 |
| **got1** | glutamic-oxaloacetic transaminase 1, soluble | 0.664 | 0.223 | 0.310 | 0.022 | 0.342 | 0.037 | 0.170 | 0.023 | 0.215 | 0.029 | -0.475 | 0.090 | -0.454 | 0.164 |
| **got2** | glutamatic-oxaloacetic transaminase 2, mitochondrial | 1.697 | 0.286 | 1.190 | 0.104 | 1.217 | 0.099 | 0.563 | 0.089 | 0.737 | 0.021 | -0.291 | 0.037 | -0.526 | 0.301 |
| **Respiratory Complexes and Related Proteins** | | | | | | | | | | | | | | | |
| **Ndufs1** | NADH dehydrogenase (ubiquinone) Fe-S protein 1 | 1.107 | 0.276 | 0.800 | 0.069 | 0.628 | 0.097 | 0.338 | 0.018 | 0.425 | 0.023 | -0.240 | -0.271 | -0.566 | 0.725 |
| **Ndufv1** | NADH dehydrogenase (ubiquinone) flavoprotein 1 | 0.888 | 0.194 | 0.690 | 0.052 | 0.585 | 0.091 | 0.325 | 0.024 | 0.341 | 0.032 | -0.188 | -0.209 | -0.522 | 0.325 |
| **sdha** | succinate dehydrogenase complex, subunit A, flavoprotein (Fp) | 2.519 | 0.254 | 2.030 | 0.248 | 2.567 | 0.129 | 0.635 | 0.034 | 0.768 | 0.065 | -0.235 | 0.343 | -0.667 | -0.102 |
| **sdhb** | succinate dehydrogenase complex, subunit B, iron sulfur (Ip) | 0.568 | 0.063 | 0.453 | 0.057 | 0.461 | 0.045 | 0.157 | 0.020 | 0.234 | 0.010 | -0.242 | 0.061 | -0.638 | 0.436 |
| **sdhc** | succinate dehydrogenase complex, subunit C, integral membrane protein | 0.220 | 0.019 | 0.201 | 0.026 | 0.265 | 0.033 | 0.100 | 0.002 | 0.099 | 0.006 | -0.138 | 0.350 | -0.466 | -0.274 |
| **etfa** | electron transferring flavoprotein, alpha polypeptide | 1.260 | 0.276 | 0.797 | 0.115 | 1.110 | 0.112 | 0.243 | 0.024 | 0.387 | 0.054 | -0.377 | 0.480 | -0.672 | 0.069 |
| **etfb** | electron transferring flavoprotein, beta polypeptide | 2.549 | 0.572 | 1.863 | 0.225 | 2.177 | 0.230 | 0.628 | 0.086 | 0.708 | 0.076 | -0.262 | 0.217 | -0.647 | -0.060 |
| **etfdh** | electron transferring flavoprotein, dehydrogenase | 0.835 | 0.197 | 0.531 | 0.064 | 0.645 | 0.057 | 0.216 | 0.018 | 0.251 | 0.010 | -0.360 | 0.271 | -0.568 | -0.079 |
| **Coq6** | coenzyme Q6 homolog (yeast) | 0.040 | 0.009 | 0.025 | 0.003 | 0.031 | 0.003 | 0.017 | 0.002 | 0.015 | 0.001 | -0.364 | 0.283 | -0.302 | -0.299 |
| **Uqcrc1** | ubiquinol-cytochrome c reductase core protein 1 | 2.984 | 0.619 | 2.244 | 0.206 | 2.157 | 0.201 | 0.625 | 0.062 | 0.865 | 0.131 | -0.234 | -0.035 | -0.715 | 0.423 |
| **Atp5a1** | ATP synthase, H+ transporting, mitochondrial F1 complex, alpha subunit 1 | 14.823 | 1.367 | 9.017 | 1.192 | 10.672 | 0.923 | 5.417 | 0.734 | 6.904 | 0.634 | -0.427 | 0.243 | -0.374 | 0.046 |
| **Atp5b** | ATP synthase, H+ transporting mitochondrial F1 complex, beta subunit | 16.525 | 1.622 | 11.896 | 1.169 | 12.496 | 0.960 | 4.973 | 0.705 | 6.771 | 0.593 | -0.298 | 0.072 | -0.578 | 0.298 |
| **Proteostasis Pathways** | | | | | | | | | | | | | | | |
| **Cryab** | crystallin, alpha B | 0.142 | 0.020 | 0.044 | 0.007 | 0.064 | 0.003 | 0.044 | 0.010 | 0.056 | 0.012 | -0.707 | 0.578 | N.S | -0.171 |
| **hsp90b1** | heat shock protein 90, beta (Grp94), member 1 | 1.056 | 0.142 | 1.293 | 0.097 | 1.112 | 0.090 | 1.236 | 0.251 | 1.243 | 0.120 | 0.157 | N.S | N.S | N.S |
| **hspa1a** | heat shock protein 1A | 1.974 | 0.110 | 1.568 | 0.137 | 1.491 | 0.119 | 1.451 | 0.224 | 1.497 | 0.295 | N.S | N.S | -0.106 | N.S |
| **hspa5** | heat shock protein 5 | 0.291 | 0.016 | 0.270 | 0.030 | 0.192 | 0.021 | 0.253 | 0.060 | 0.291 | 0.034 | -0.139 | -0.253 | N.S | 0.542 |
| **hspa9** | heat shock protein 9 | 0.785 | 0.163 | 0.611 | 0.102 | 0.561 | 0.069 | 0.217 | 0.045 | 0.235 | 0.020 | -0.160 | N.S | -0.605 | N.S |
| **hspd1** | heat shock protein 1 (chaperonin) | 1.407 | 0.194 | 1.259 | 0.108 | 1.023 | 0.102 | 0.687 | 0.115 | 0.792 | 0.016 | N.S | -0.214 | -0.479 | 0.532 |
| **lonp1** | lon peptidase 1, mitochondrial | 0.205 | 0.046 | 0.176 | 0.015 | 0.204 | 0.011 | 0.095 | 0.006 | 0.117 | 0.007 | N.S | 0.168 | -0.449 | N.S |
| **lonp2** | lon peptidase 2, peroxisomal | 0.023 | 0.003 | 0.021 | 0.004 | 0.024 | 0.002 | 0.008 | 0.003 | 0.012 | 0.004 | -0.205 | 0.332 | -0.625 | 0.175 |
| **Peroxisomal Proteins** | | | | | | | | | | | | | | | |
| **Abcd3** | ATP-binding cassette, sub-family D (ALD), member 3 | 0.465 | 0.058 | 0.544 | 0.071 | 0.645 | 0.090 | 0.217 | 0.022 | 0.200 | 0.015 | N.S | 0.262 | -0.560 | -0.262 |
| **Acox1** | acyl-Coenzyme A oxidase 1, palmitoyl | 0.327 | 0.063 | 0.429 | 0.071 | 0.487 | 0.075 | 0.087 | 0.008 | 0.142 | 0.040 | N.S | 0.278 | -0.752 | 0.187 |
| **Ech1** | enoyl coenzyme A hydratase 1, peroxisomal | 0.927 | 0.127 | 0.917 | 0.118 | 1.104 | 0.101 | 0.366 | 0.031 | 0.392 | 0.016 | N.S | 0.247 | -0.580 | -0.134 |
| **Ephx2** | Epoxide Hydrolase 2 | 0.783 | 0.141 | 1.790 | 0.357 | 2.004 | 0.342 | 0.317 | 0.110 | 0.326 | 0.016 | 0.820 | 0.289 | -0.809 | N.S |
| **mdh1** | malate dehydrogenase 1, NAD (soluble) | 8.040 | 0.796 | 6.655 | 0.859 | 7.770 | 0.741 | 2.961 | 0.234 | 3.708 | 0.597 | -0.218 | 0.221 | -0.525 | N.S |
| **Pecr** | peroxisomal trans-2-enoyl-CoA reductase | 2.617 | 0.911 | 3.403 | 0.702 | 3.605 | 0.611 | 0.254 | 0.086 | 0.394 | 0.128 | 0.269 | 0.276 | -0.918 | 0.333 |
| **slc25a20** | solute carrier family 25 (mitochondrial carnitine/acylcarnitine translocase) 20 | 0.485 | 0.090 | 0.330 | 0.027 | 0.274 | 0.025 | 0.195 | 0.018 | 0.220 | 0.020 | -0.305 | -0.170 | -0.400 | 0.363 |
| **Other Mitochondrial Proteins** | | | | | | | | | | | | | | | |
| **Atp2a2** | ATPase, Ca++ transporting, cardiac muscle, slow twitch 2 | 0.139 | 0.010 | 0.138 | 0.009 | 0.159 | 0.035 | 0.117 | 0.003 | 0.179 | 0.018 | N.S | 0.046 | -0.139 | 0.443 |
| **cd36** | CD36 antigen | 0.151 | 0.019 | 0.159 | 0.026 | 0.136 | 0.024 | 0.034 | 0.005 | 0.041 | 0.007 | -0.071 | -0.106 | -0.761 | 0.345 |
| **Clpp** | caseinolytic mitochondrial matrix peptidase proteolytic subunit | 0.214 | 0.047 | 0.266 | 0.029 | 0.283 | 0.036 | 0.121 | 0.020 | 0.133 | 0.003 | 0.269 | 0.077 | -0.537 | 0.059 |
| **Clpx** | caseinolytic mitochondrial matrix peptidase chaperone subunit | 0.105 | 0.013 | 0.121 | 0.011 | 0.122 | 0.008 | 0.127 | 0.005 | 0.146 | 0.007 | 0.137 | 0.035 | 0.089 | 0.109 |
| **ckmt1** | creatine kinase, mitochondrial 2 | 1.898 | 1.389 | 0.473 | 0.079 | 0.568 | 0.091 | 0.314 | 0.148 | 0.245 | 0.068 | -0.543 | 0.227 | -0.460 | -0.100 |
| **prkaca** | protein kinase, cAMP dependent, catalytic, alpha | 0.069 | 0.006 | 0.052 | 0.003 | 0.050 | 0.003 | 0.059 | 0.003 | 0.061 | 0.002 | -0.239 | -0.032 | 0.133 | 0.076 |
| **Rhot1** | Ras Homolog Family Member T1 | 0.162 | 0.012 | 0.144 | 0.012 | 0.159 | 0.010 | 0.076 | 0.002 | 0.096 | 0.003 | -0.129 | 0.127 | -0.459 | 0.130 |
| **Samm50** | sorting and assembly machinery component 50 homolog (S. cerevisiae) | 0.395 | 0.039 | 0.275 | 0.016 | 0.248 | 0.036 | 0.167 | 0.018 | 0.209 | 0.011 | -0.305 | -0.149 | -0.394 | 0.491 |
| **slc25a11** | solute carrier family 25 (mitochondrial carrier oxoglutarate carrier), member 11 | 0.925 | 0.231 | 0.662 | 0.069 | 0.627 | 0.050 | 0.297 | 0.017 | 0.364 | 0.022 | -0.266 | N.S | -0.529 | 0.229 |
| **Slc25a4** | solute carrier family 25 (mitochondrial carrier, adenine nucleotide translocator), member | 1.932 | 0.029 | 1.348 | 0.091 | 1.914 | 0.302 | 0.909 | 0.038 | 1.201 | 0.074 | -0.313 | 0.348 | -0.324 | N.S |
| **Slc25a4/5/31** | solute carrier family 25 (mitochondrial carrier, adenine nucleotide translocator), member | 41.064 | 3.015 | 33.056 | 2.755 | 38.219 | 2.826 | 13.184 | 0.955 | 18.025 | 2.460 | -0.211 | 0.171 | -0.592 | 0.153 |
| **Tufm** | Tu translation elongation factor, mitochondrial | 0.412 | 0.079 | 0.357 | 0.031 | 0.360 | 0.024 | 0.132 | 0.010 | 0.192 | 0.019 | -0.118 | 0.024 | -0.622 | 0.416 |

*** The magnitude of effects are calculated only for those with significant changes; NS= not significant**
